# Comparative genomics of human *Lactobacillus crispatus* isolates reveals genes for glycosylation and glycogen degradation: Implications for *in vivo* dominance of the vaginal microbiota

**DOI:** 10.1101/441972

**Authors:** Charlotte van der Veer, Rosanne Y. Hertzberger, Sylvia M. Bruisten, Hanne L. P. Tytgat, Jorne Swanenburg, Alie de Kat Angelino-Bart, Frank Schuren, Douwe Molenaar, Gregor Reid, Henry de Vries, Remco Kort

**Author notes:** Corresponding author at Netherlands Organization for Applied Scientific Research (TNO), Microbiology and Systems Biology, Utrechtseweg 48, 3704 HE, Zeist, the Netherlands.

## Abstract

**Background:** A vaginal microbiota dominated by lactobacilli (particularly *Lactobacillus crispatus*) is associated with vaginal health, whereas a vaginal microbiota not dominated by lactobacilli is considered dysbiotic. Here we investigated whether *L. crispatus* strains isolated from the vaginal tract of women with *Lactobacillus*-dominated vaginal microbiota (LVM) are pheno- or genotypically distinct from *L. crispatus* strains isolated from vaginal samples with dysbiotic vaginal microbiota (DVM).

**Results:** We studied 33 L. crispatus strains (n=16 from LVM; n=17 from DVM). Comparison of these two groups of strains showed that, although strain differences existed, both groups were heterofermentative, produced similar amounts of organic acids, inhibited *Neisseria gonorrhoeae* growth and did not produce biofilms. Comparative genomics analyses of 28 strains (n=12 LVM; n=16 DVM) revealed a novel, 3-fragmented glycosyltransferase gene that was more prevalent among strains isolated from DVM. Most *L. crispatus* strains showed growth on glycogen-supplemented growth media. Strains that showed less efficient (n=6) or no (n=1) growth on glycogen all carried N-terminal deletions (respectively, 29 and 37 amino acid-deletions) in a putative pullulanase type I gene.

**Discussion:** *L. crispatus* strains isolated from LVM were not phenotypically distinct from *L. crispatus* strains isolated from DVM, however, the finding that the latter were more likely to carry a 3-fragmented glycosyltransferase gene may indicate a role for cell surface glycoconjugates, which may shape vaginal microbiota-host interactions. Furthermore, the observation that variation in the pullulanase type I gene associated with growth on glycogen discourages previous claims that *L. crispatus* cannot directly utilize glycogen.

## INTRODUCTION

The vaginal mucosa hosts a community of commensal, symbiotic and sometimes pathogenic micro-organisms. Increasing evidence has shown that the bacteria within this community, referred to here as the vaginal microbiota (VM), play an important role in protecting the vaginal tract from pathogenic infection, which can have far reaching effects on a woman’s sexual and reproductive health [1, 2]. Several VM compositions have been described, including VM dominated by: 1) *Lactobacillus iners;* 2) *L. crispatus;* 3) *L. gasseri*; 4) *L. jensenii* and; 5) VM that are not dominated by a single bacterial species but rather consist of diverse anaerobic bacteria, including *Gardnerella vaginalis* and members of Lachnospiraceae and Leptotrichiaceaeprevotella [3-5]. Particularly VM that are dominated by *L. crispatus* are associated with vaginal health, whereas a VM consisting of diverse anaerobes – commonly referred to as vaginal dysbiosis - have been shown to increase a woman’s odds for developing bacterial vaginosis (BV), acquiring STI’s, including HIV, and having an adverse pregnancy outcome [1, 2, 4, 6].

The application of human vaginal *L. crispatus* isolates as therapeutic agents to treat dysbiosis may have much potential [7, 8], but currently there are still many gaps in our knowledge concerning the importance of specific physiological properties of *L. crispatus* for a sustained domination on the mucosal surface of the vagina. Comparative genomics approaches offer a powerful tool to identify novel important physiological properties of bacterial strains. The genomes of nine human *L. crispatus* isolates have previously been studied, also in the context of vaginal dysbiosis [9, 10]. Comparative genomics of these strains showed that about 60% of orthologous groups (genes derived from the same ancestral gene) were conserved among all strains; i.e. comprising a ‘core’ genome [10]. The accessory genome was defined as genes shared by at least two strains, while unique genes are specific to a single strain. Currently it is unclear whether traits pertaining to *in vivo* dominance are shared by all strains (core genome), or only by a subset of strains (accessory genome). For example, both women with and without vaginal dysbiosis can be colonized with *L. crispatus* (see e.g.[11]) and we do not yet fully understand why in some women *L. crispatus* dominates and in others not.

The following bacterial traits may be of importance for *L. crispatus* to successfully dominate the vaginal mucosa: 1) the formation of an extracellular matrix (biofilm) on the vaginal mucosal surface; 2) the production of antimicrobials such as lactic acid, bacteriocins and H_2_O_2_ that inhibit the growth and/or adhesion of urogenital pathogens; 3) efficient utilization of available nutrients – particularly glycogen, as this is the main carbon source in the vaginal lumen; and; 4) the modulation of host-immunogenic responses. Considering these points, firstly, Ojala *et al*. [10] observed genomic islands encoding enzymes involved in exopolysacharide (EPS) biosynthesis in the accessory genome of *L. crispatus* and postulated that strain differences in this trait could contribute to differences in biofilm formation, adhesion and competitive exclusion of pathogens. Secondly, experiments have shown that *L. crispatus* effectively inhibits urogenital pathogens through lactic acid production, but these studies included only strains originating from healthy women [12-16]. Abdelmaksoud *et al*. [9] compared *L. crispatus* strains isolated from *Lactobacillus*-dominated VM (LVM) with strains isolated from dysbiotic VM (DVM) and indeed observed decreased lactic acid production in one of the strains isolated from DVM, providing an explanation for its low abundance. However, no significant conclusion could be made as their study included only eight strains. Thirdly, there is a general consensus that vaginal lactobacilli (including *L. crispatus*) ferment glycogen thus producing lactic acid, but no actual evidence exists that *L. crispatus* produces the enzymes to directly degrade glycogen [10, 17]. Lastly, *L. crispatus*-dominated VM are associated with an anti-inflammatory vaginal cytokine profile [18, 19] and immune evasion is likely a crucial (but poorly studied) factor that allows *L. crispatus* to dominate the vaginal niche. A proposed underlying mechanism is that *L. crispatus* produces immunomodulatory molecules [20], but *L. crispatus* may also accomplish immune modulation by alternating its cell surface glycosylation, as has been suggested for gut commensals [21]. Taken together, there is a clear need to study the properties of more human (clinical) *L. crispatus* isolates to fully appreciate the diversity within this species.

Here we investigated whether *L. crispatus* strains isolated from the vaginal tract of women with LVM are pheno- or genotypically distinct from *L. crispatus* strains isolated from vaginal samples with DVM, with the aim to identify bacterial traits pertaining to a successful domination of lactobacilli of the vaginal mucosa.

## RESULTS

### Lactobacillus crispatus strain selection and whole genome sequencing

For this study, 40 nurse-collected vaginal swabs were obtained from the Sexually Transmitted Infections clinic in Amsterdam, the Netherlands, from June to August 2012, as described previously by Dols *et al*. [4]. In total, 33 *L. crispatus* strains were isolated from these samples (n=16 from LVM samples; n=17 *L. crispatus* strains from DVM samples). Following whole genome sequencing, four contigs (n=3 strains from LVM; n=1 strains from DVM) were discarded as they had less than 50% coverage with other assemblies or with the reference genome (ST1), suggesting that these isolates belonged to a different *Lactobacillus* species. One contig (from a strain isolated from LVM) aligned to the reference genome, but its genome size was above the expected range, suggestive of contamination with a second strain and was therefore also discarded. The remaining 28 isolates (n=12 LVM and n=16 DVM) were assembled and used for comparative genomics. These genomes have been deposited at DDBJ/ENA/GenBank under the accession numbers NKKQ00000000-NKLR00000000. The versions described in this paper are versions NKKQ01000000-NKLR01000000 (Table 1).

**Table 1.**
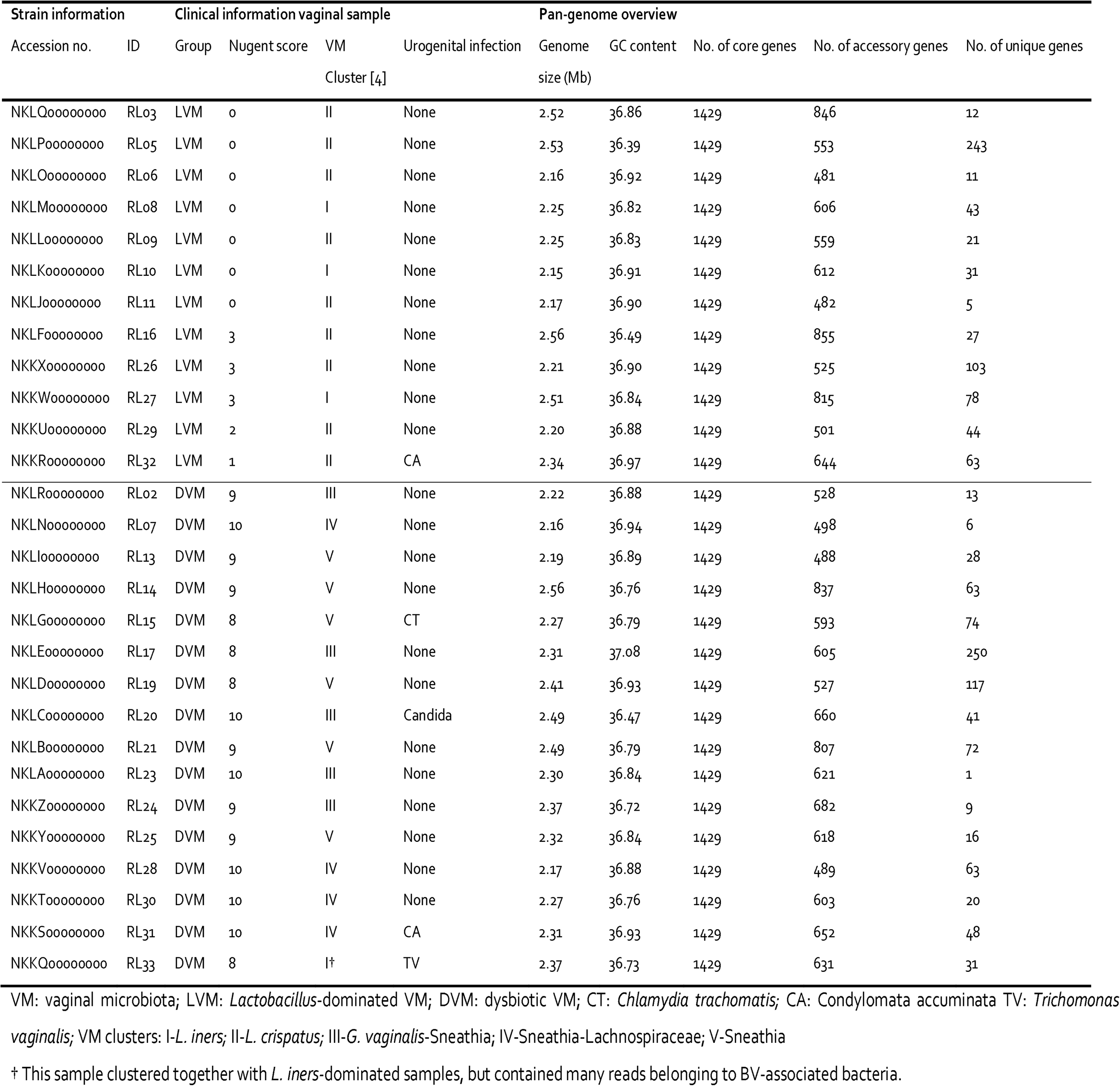
Overview and properties of 28 *L. crispatus* strains isolated from vaginal swabs with *Lactobacillus*-dominated vaginal microbiota or dysbiotic vaginal microbiota.

### Lactobacillus crispatus pan genome

The 28 *L. crispatus* genomes had an average length of 2.31 Mbp (range 2.16 – 2.56 MB) (Table 1), which was slightly larger than the reference genome (ST1; 2.04Mbp). The GC content of the genomes was on average 36.8%, similar to other lactobacilli [10]. An average of 2099 genes were annotated per strain (Table 1; Figure 1). This set of 28 *L. crispatus* genomes comprised 4261 different gene families. The core genome consisted of 1429 genes (which corresponds to ~68% of a given genome) and the accessory genome averaged at 618 genes (~30%) per strain. Each strain had on average 54 unique genes (~2.0%). The number of accessory and unique genes did not significantly differ between strains isolated from LVM or from DVM, with respectively an average of 621 (range: 481-855) and 55 (range: 5-243) genes for LVM strains and 615 (range: 488-837) and 53 (range: 1-250) genes for DVM strains. The distribution of cluster of ortholog groups (COG) also did not differ between strains from *Lactobacillus*-dominated and DVM. The gene accumulation model [22] describes the expansion of the pan-genome as function of the number of genomes and estimated that this species has access to a larger gene pool than described here; the model estimated the *L. crispatus* pan genome to include 4384 genes.

**Figure 1.**
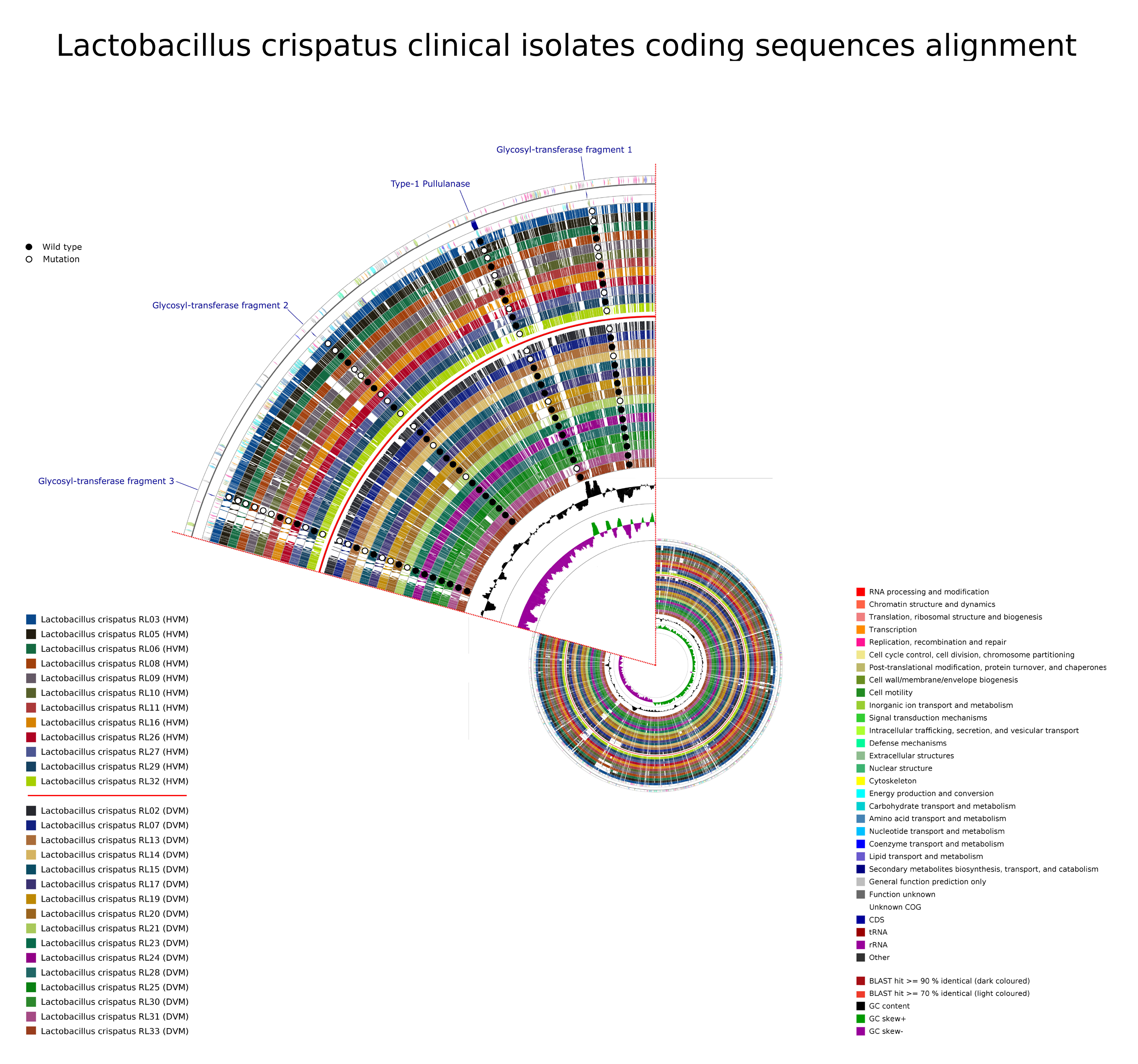
Whole genome alignments of the coding sequences from the *Lactobacillus crispatus* clinical isolates described in this study. The outermost ring represents COG annotated genes on the forward strand (color coded according to the respective COG). The positions of the genes discussed in this article are indicated. The third ring represents COG annotated genes on the reverse strand (color coded according to the respective COG). The next twelve rings each represent one genome of the LVM strains, followed by a separator ring and 16 rings each representing a genome of the DVM strains. The height of the bar and the saturation of the color in these rings indicate a BLAST hit of either >90% identity (darker colored) or >70% identity (lightly colored). Hits below 70% identity score are not shown and appear as white bars in the plots. The two inner most rings represent the GC content of that area and the GC-skew respectively. The presence or absence of the gene variants discussed in this article is indicated in each genome by black and white dots. A black dot indicates that a wild-type gene (as compared to the STI reference genome) is present in that genome, a white dot indicates that no copy of that gene (fragment) was present or that it carried a deletion (for the type 1 pullulanase). Abbreviations: COG: cluster ortholog genes; LVM: *Lactobacillus*-dominated vaginal microbiota; DVM: dysbiotic vaginal microbiota; WT: wild type.

### A fragmented glycosyltransferase gene was abundant among strains isolated from DVM

In a comparative genomics analysis we aimed to identify genes that were specific to strains isolated from either LVM or DVM. We observed that three transposases, one of which was further classified as an IS30 family transposase, were more abundant among strains isolated from DVM than among strains from LVM. IS30 transposases are associated with genomic instability and have previously been found to flank genomic deletions in commercial *L. rhamnosus* GG probiotic strains [23]. Most notably, we observed that strains from DVM were more likely to carry three gene fragments of a single glycosyltransferase (GT) than strains isolated from LVM. GTs are enzymes that are involved in the transfer of a sugar moiety to a substrate and are thus essential in synthesis of glycoconjugates like exopolysaccharides, glycoproteins and glycosylated teichoic acids [24, 25].

The three differentially abundant GT gene fragments all align to different regions of a family 2 A-fold GT of the ST1 *L. crispatus* strain (CGA_000165885.1) and are flanked by other genes potentially encoding GTs (Figure 2). Fragment 1 aligns with 472 bp of the original unfragmented GT, while fragment 2 overlaps with the last 3 bp of fragment 1 and fragment 3 overlaps 7 bp with fragment 2. Given that all these fragments align to the non-fragmented GT gene in in *L. crispatus* ST1, we hypothesize that the three fragments belong to the same GT. The *L. crispatus* genomes however contained a combination of one or more of the three GT fragments, while the surrounding genes were conserved among the strains. The first fragment of 510 bp contains the true GT fold domain and is thus responsible for the catalytic activity of the GT. The second and third fragment are considerably shorter, respectively 228 and 328 bp, and do not harbor any significant relation to a known GT-fold (Figure 3). Four different combinations of GT fragments were observed in the studied genomes, namely a variant with: (1) no fragments, (2) all three fragments, (3) fragment 1 and 3, and (4) fragment 1 and 2 (Figure 2; Table 2).

**Figure 2.**
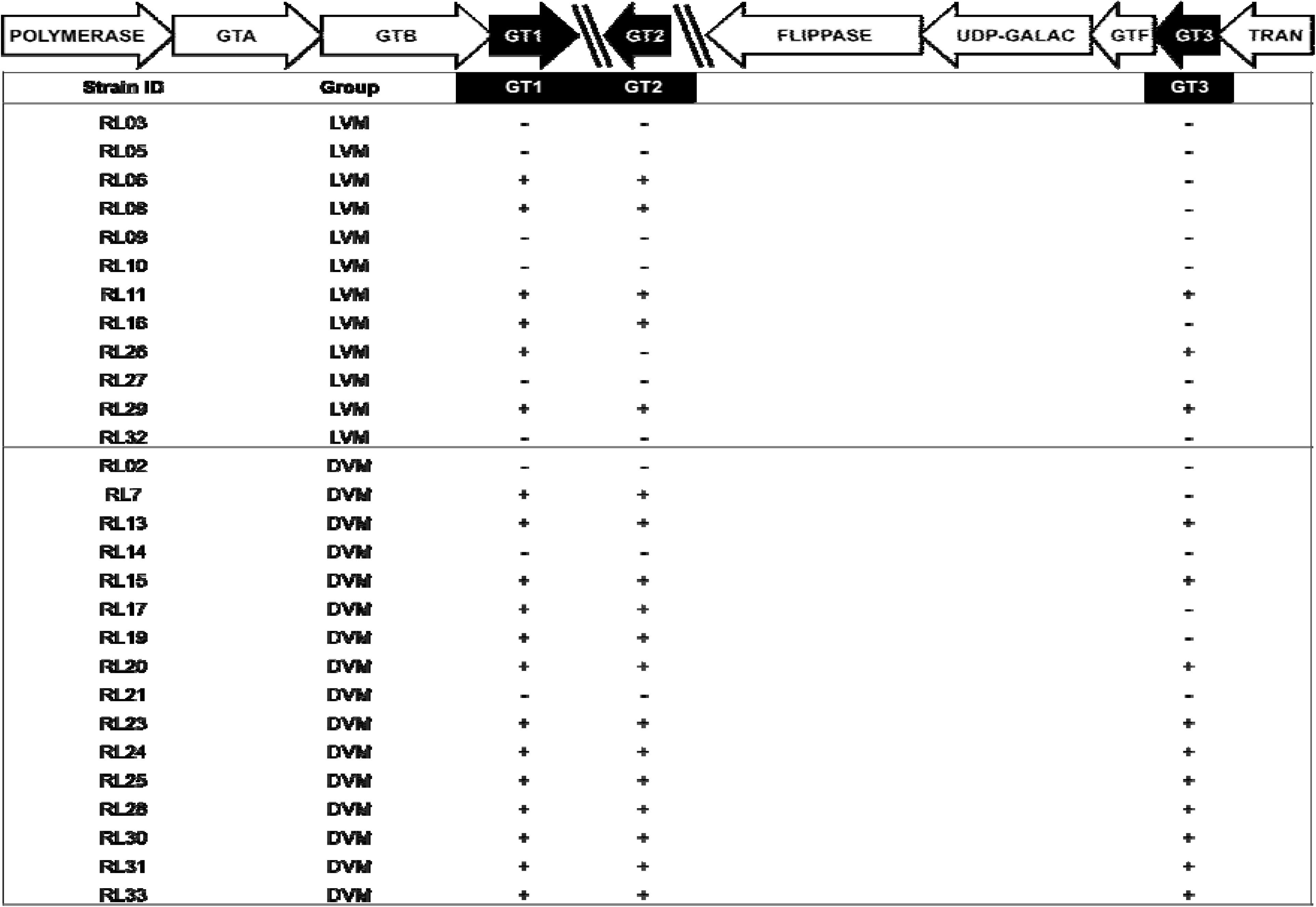
Schematic overview of the organization of the glycosyltransferase fragments in the *Lactobacillus crispatus* genomes. The orientation of the fragments is dependent on the assembly, and can therefore be different than depicted here. Also, the distance between the fragments is undetermined and can be of any length (depicted with diagonal lines). Abbreviations: GT: Glycosyltransferase; GTA, GTB: GT super families; GT1, GT2, GT3: GT fragments 1, 2, 3; UDP-GALAC: UDP-Galactopyranose mutase; GTF: GT family 1; TRAN: transposase; LVM: *Lactobacillus*-dominated vaginal microbiota; DVM: dysbiotic vaginal microbiota.

**Figure 3.**
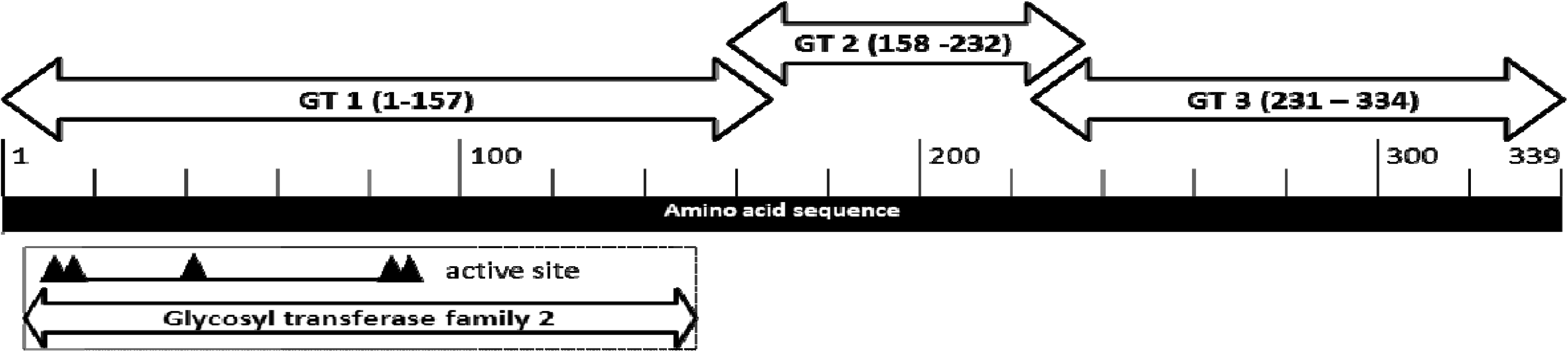
Schematic overview of how the glycosyltransferase fragments align to the *Lactobacillus crispatus* ST1 reference genome. The first fragment comprises the conserved glycosyltransferase family 2 domain with catalytic activity. The shorter second and third fragments most probably do not harbor any catalytic GT activity. We hypothesize that these two fragments play a role in steering the specific activity of the GT (e.g. towards donor or substrate specificity). Abbreviation: GT: glycosyltransferase.

**Table 2.** Comparison of distribution of glycosyltransferase (GT) gene fragments in *Lactobacillus crispatus* genomes isolated from vaginal samples with *Lactobacillus*-dominated or dysbiotic vaginal microbiota.

|  | **LVM** | **DVM** | **p-value\*** |
| --- | --- | --- | --- |
|  | **N = 12 (%)** | **N = 16 (%)** |  |
| No GT fragments | 6 (50.0) | 3 (18.8) | 0.114 |
| 1<sup>st</sup> and 2<sup>nd</sup> GT fragments | 3 (25.0) | 3 (18.8) | 1.000 |
| 1<sup>st</sup> and 3<sup>rd</sup> GT fragment | 1 (8.3) | 0 (0.0) | 0.429 |
| All 3 GT fragments | 2 (16.6) | 10 (62.5) | **0.023** |

LVM: *Lactobacillus*-dominated VM; DVM: dysbiotic VM
\* Fisher's Exact test.

### Strains isolated from LVM were not phenotypically distinct from strains isolated from DVM

Phenotypic studies on the *L. crispatus* strains did not reveal any biofilm formation – as assessed by crystal violet assays, except for one strain (RL19) which produced a weak biofilm. In line with this, very low levels of autoaggregation (on average 5%) were observed and this also did not differ between the two groups of strains. Strain specific carbohydrate fermentation profiles were observed, as assessed by a commercial API CH50 test, but the distribution of these profiles did not relate to whether the strains were isolated from LVM or from DVM. Strains isolated from LVM produced similar amounts of organic acids compared with strains isolated from DVM when grown on chemically defined medium mimicking vaginal fluids [26]. The strains mainly produced lactic acid. Other acids such as succinate acid, butyric acid, glutamic acid, phenylalanine, isoleucine and tyrosine were also produced, but four-fold lower compared to lactic acid. Very small acidic molecules, such as acetic and propionic acid, were out of the detection range and could thus not be measured. We also assessed antimicrobial activity against a common urogenital pathogen *Neisseria gonorrhoeae*. Inhibition was similar for strains isolated from LVM and from DVM: *N. gonorrhoeae* growth was inhibited (i.e. lower OD_600nm_ in stationary phase compared to the control), in a dose-dependent way, by on average 27.9 ± 15.8% for undiluted *L. crispatus* supernatants compared to the *N. gonorrhoeae* control. Undiluted neutralized *L. crispatus* supernatants inhibited *N. gonorrhoeae* growth by on average 15.7 ± 16.3% (Supplementary information).

### Strain-specific glycogen growth among both LVM and DVM isolates

Of the 28 strains for which full genomes were available, we tested 25 strains (n=12 LVM and n=13 DVM) for growth on glycogen. We compared growth on glucose-free NYCIII medium supplemented with glycogen as carbon source to growth on NYCIII medium supplemented with glucose (positive control) and NYCIII medium supplemented with water (negative control). All except one strain (RL05) showed growth on glycogen; however six strains showed substantially less efficient growth on glycogen. One strain showed a longer lag time (RL19; on average 4.5 hours, compared to an average of 1.5 hours for other strains) and five strains (RL02, RL06, RL07, RL09 and RL26) showed a lower OD after 36 hours of growth compared to other strains (Figure 4). Growth on glycogen did not correlate to whether the strain was isolated from LVM or DVM.

**Figure 4.**
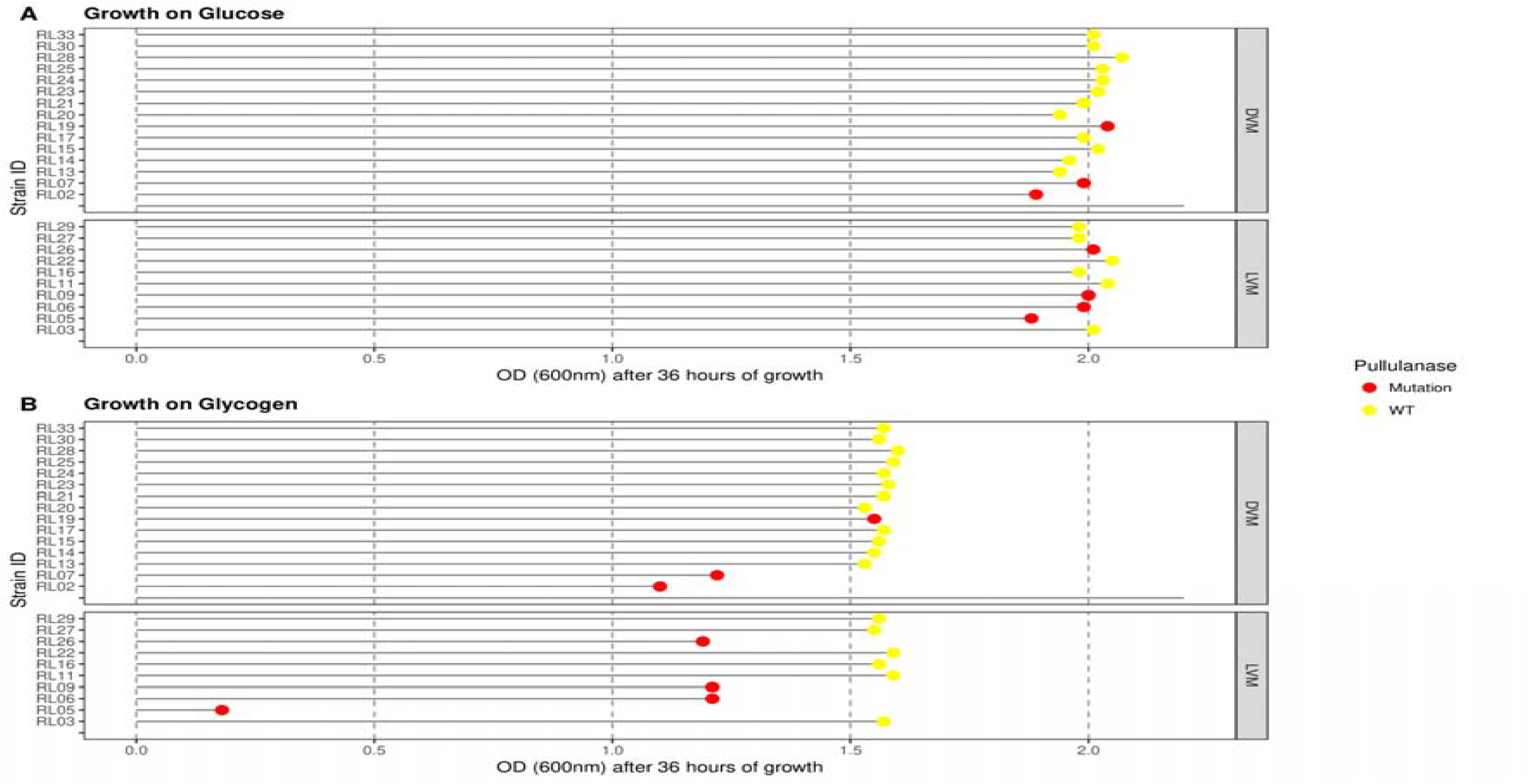
Growth on glycogen for *Lactobacillus crispatus* strains isolated from *Lactobacillus*-dominated and from dysbiotic vaginal microbiota. Strains were grown in minimal medium supplemented with A) 5% glucose and B) 5% glycogen. Strains that showed less efficient or no growth on glycogen carried a mutation in the N-terminal sequence of a putative type I pullulanase gene. RL19 showed a longer lag time compared to other strains; on average 4.5 hours, compared to an average of 1.5 hours for other strains. Abbreviations: LVM: *Lactobacillus*-dominated vaginal microbiota; DVM: dysbiotic vaginal microbiota; WT: wild type.

### Growth on glycogen corresponded with variation in a putative pullulanase type I gene

We followed-up on the glycogen growth experiments with a gene-trait analysis as glycogen is considered to be a key, although disputed, nutrient (directly) available to *L. crispatus*. We searched the *L. crispatus* genomes for the presence/absence of enzymes that can potentially be involved in glycogen metabolism. We thus searched for orthologs of the: 1) glycogen debranching enzyme (encoded by *glgX*) in *Escherichia coli* [27, 28]; 2) *Streptococcus agalactiae* pullulanase [29]; 3) SusB of *Bacteroides thetaiotaomicron* [30]; and 4) the amylase (encoded by *amyE*) of *Bacillus subtilis* [31]. This search revealed a gene that was similar to the *glgX* gene; this gene was annotated as a pullulanase type I gene. In other species this pullulanase is bound to the outer S-layer of the cell wall, suggesting that this enzyme utilizes extracellular glycogen [32]. All except two strains (RL31, RL32) carried a copy of this gene. The genes are conserved except for variation in the N-terminal sequence that encodes a putative signal peptide that may be involved in subcellular localization of the enzyme. All strains with less efficient growth on glycogen had a 29 amino acid deletion in the N-terminal sequence (strains: RL02, RL06, RL07, RL09, RL19 and RL26) and the strain that showed no growth (RL05) had an 8 amino acid deletion in the same region as the other strains in addition to 37 amino acid deletion further downstream (Table 3).

**Table 3.**
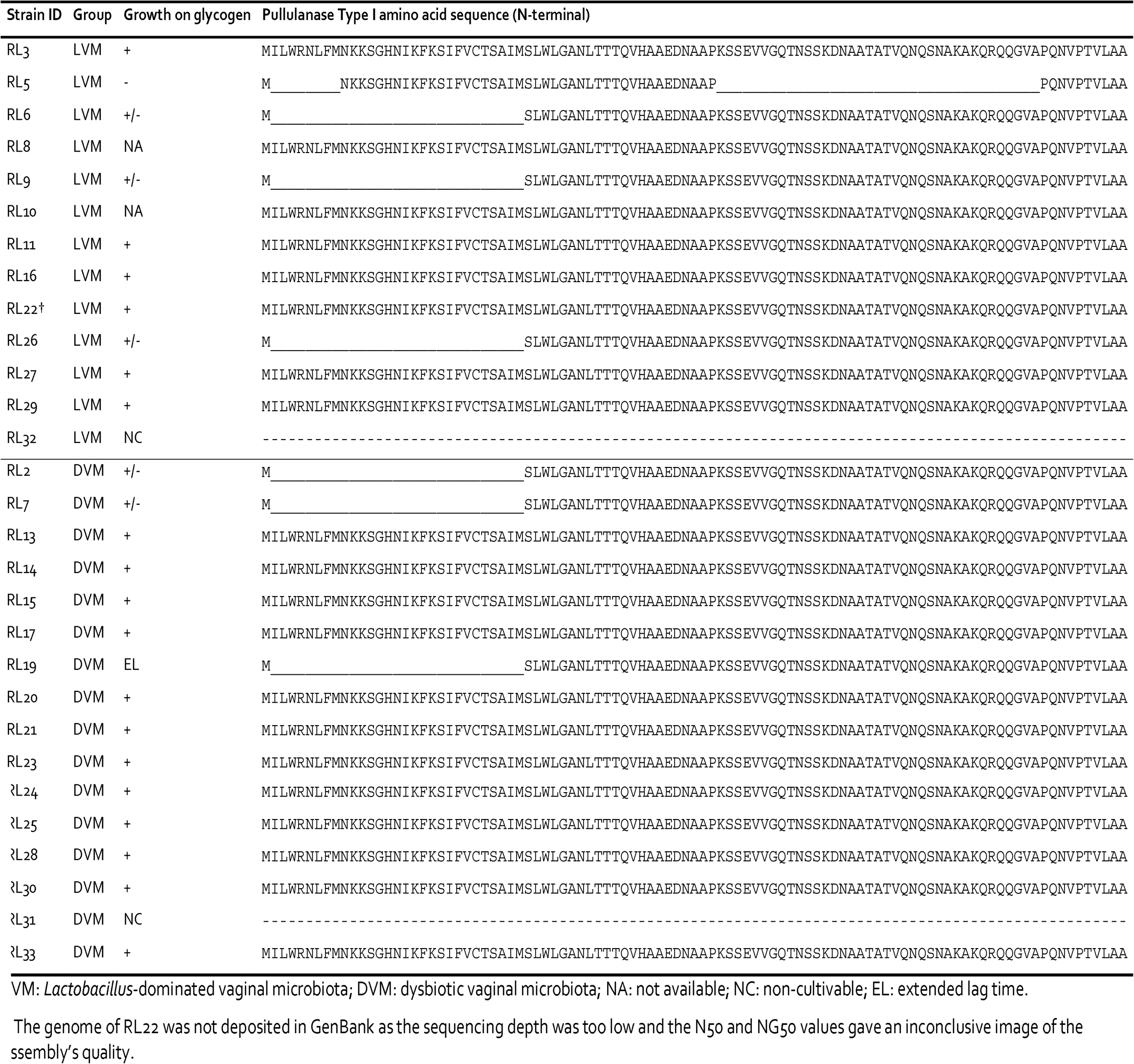
Overview of *Lactobacillus crispatus* strain specific growth on glycogen and corresponding translated amino acid sequence at the N-terminal of a lulanase type I gene.

## DISCUSSION

### Key findings of this paper

Here we report the full genomes of 28 *L. crispatus* clinical isolates; the largest contribution of *L. crispatus* clinical isolates to date. These strains were isolated from women with LVM and from women with DVM. A comparative genomics analysis revealed that a glycosyltransferase gene was more frequently found in the genomes of strains isolated from DVM as compared with strains isolated from LVM, suggesting a fitness advantage for carrying this gene in *L. crispatus* under dysbiotic conditions and a role of surface glycoconjugates in microbiota-host interactions. Comparative experiments pertaining to biofilm formation, antimicrobial activity and nutrient utilization showed that these two groups of strains did not phenotypically differ from each other. Of particular novelty value, we found that these clinical *L. crispatus* isolates were capable of growth on glycogen and that variation in a pullulanase type I gene correlates to the level of this activity.

### Vaginal dysbiotic conditions may pressurize *Lactobacillus crispatus* to vary its glycome

Several studies have shown that vaginal dysbiosis is associated with an increased pro-inflammatory response, including an increase in pro-inflammatory chemokines and cytokines, but also elevated numbers of activated CD4+ T cells [3, 19], although no clinical signs of inflammation are present and vaginal dysbiosis is seen as a condition rather than as a disease [33]. Nonetheless, it indicates that the vaginal niche in a dysbiotic state is indeed under some immune pressure and that immune evasion could be a key (but poorly studied) trait for probiotic bacterial survival and dominance on the vaginal mucosa.

Our comparative genomics analysis revealed a glycosyltransferase gene (GT) gene that was more common in strains isolated from DVM compared with strains isolated from LVM. The identified GT consists of three fragments, which all align to a single GT in the reference *L. crispatus* genome (ST1). Sequence analyses showed that the first and longest fragment exhibits close homology to a known GT-A fold and most probably harbors the active site of the GT (Figure 3). The latter two fragments do not harbor any structural motifs resembling known GTs and most probably do not harbor any catalytic GT activity. We hypothesize that these two fragments play a role in steering the specific activity of the GT (e.g. towards donor or substrate specificity). This might point towards *L. crispatus* harnessing its genetic potential to change its surface glycome. Such a process is termed phase variation and allows bacteria to rapidly adapt and diversify their surface glycans, resulting in an evolutionary advantage in the arms race between the immune system and invading bacteria. Modulation of the surface glycome by phase variation of the GT coding sequence is a common immune evasion strategy, which has been extensively studied in pathogenic bacteria like *Campylobacter jejuni* [25], but could be utilized by commensals as well [21]. We hypothesize that *L. crispatus* in DVM exploits this genetic variation to allow for (a higher) variation in cell wall glycoconjugates providing a mechanism for *L. crispatus* to persist at low levels in DVM and remain stealth from the immune system (Figure 5). Of note, evidence for expression of all of the 3 GT-fragments comes from a recent transcriptomics study that studied the effect of metronidazole treatment on the VM of women with (recurring) BV [11]. Personal communication with Dr. Zhi-Luo Deng revealed that high levels of expression for the three putative GT peptides were present in the vaginal samples of two women who were responsive to treatment (i.e. their VM was fully restored to a *L. crispatus*-dominated VM following treatment). This finding is in line with our hypothesis that the presence of the fragmented GT gene has a selective advantage for *L. crispatus* under dysbiotic conditions. Further functional experiments are needed to test this hypothesized host-microbe interaction and to coin if and how the variation of glycoconjugates is affected by this GT. Additionally, the immunological response of the host must be further studied in reference to these hypothesized microbial adaptations. The bacterial surface glycome and related variability events are currently overlooked features in probiotic strain selection, while they might be crucial to a strain’s survival and *in vivo* dominance [21].

**Figure 5.**
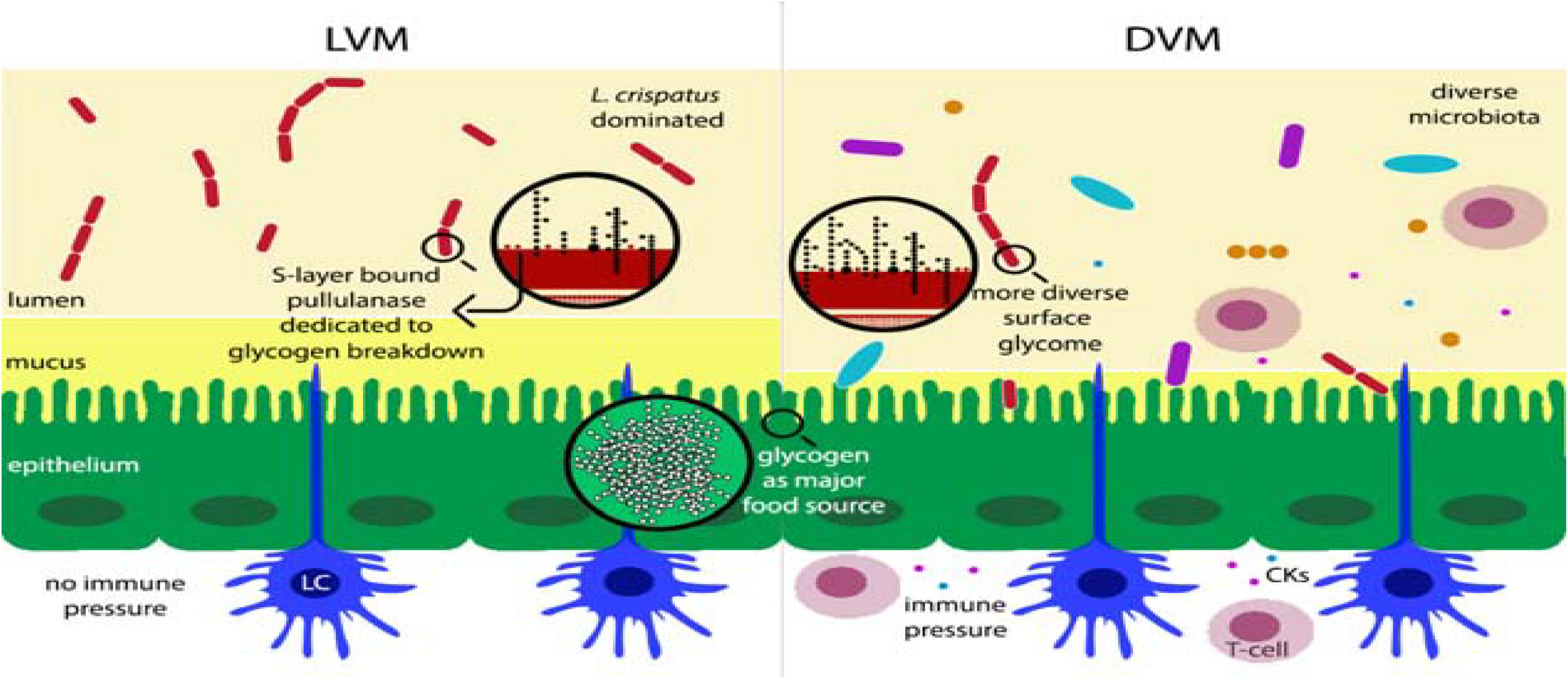
Model for enzymatic activity in glycosylation and glycogen degradation in *Lactobacillus crispatus*. Schematic representation of the vaginal environment with either LVM or DVM. Our comparative genomics analysis revealed a glycosyltransferase gene that was more common in *Lactobacillus crispatus* strains isolated from LVM (red bacteria) and DVM (low abundance of red lactobacilli, diverse bacterial population in multiple colors and forms, thinner mucus layer). We hypothesize that *L. crispatus* in DVM exploits this genetic variation to allow for (a higher) variation in cell wall glycoconjugates providing a mechanism for *L. crispatus* to persist at low levels in DVM and remain stealth from the immune system. Another finding of this work describes the ability of *L. crispatus* strains to utilize glycogen as a food source, which is associated with the presence of a full-length pullulanase gene (red dots on cell wall of *L. crispatus*). Abbreviations: LVM: *Lactobacillus*-dominated vaginal microbiota; DVM: dysbiotic vaginal microbiota, LC: Langerhans cell, CK: cytokines.

### No distinct phenotypes pertaining to dominance *in vivo* were observed

It has previously been postulated, relying merely on genomics data, that the accessory genome of *L. crispatus* could lead to strain differences relating to biofilm formation, adhesion and competitive exclusion of pathogens [9, 10]; all of which could influence whether a strain dominates the vaginal mucosa or not. Our comparative experimental work, however, showed that *L. crispatus* - irrespective of whether the strain was isolated from a woman with LVM or with DVM – all formed little to no biofilm, demonstrated effective lactic acid production and effective antimicrobial activity against *N. gonorrhoeae*. The previous genomic analyses also suggested that *L. crispatus* is herterofermentative [10]. Indeed, we observed that *L. crispatus* ferments a broad range of carbohydrates, as assessed by a commercial API test, but these profiles did not differ between strains isolated from LVM or from DVM.

### First evidence showing that *Lactobacillus crispatus* grows on glycogen

The vaginal environment of healthy reproductive-age women is distinct from other mammals in that it has low microbial diversity, a high abundance of lactobacilli and high levels of lactic acid and luminal glycogen [34]. It has been postulated that proliferation of vaginal lactobacilli is supported by estrogen-driven glycogen production [35], however the ‘fly in the ointment’ - as finely formulated by Nunn *et al*. [17] - is that evidence for direct utilization of glycogen by vaginal lactobacilli is absent. Moreover, previous reports have stated that the core genome of *L. crispatus* does not contain the necessary enzymes to break down glycogen [10, 36]. It has even been suggested that *L. crispatus* relies on amylase secretion by the host or other microbes for glycogen breakdown [17, 37], as *L. crispatus* does contain all the appropriate enzymes to consume glycogen breakdown products such as glucose and maltose [36].

Here we provide the first evidence suggesting that *L. crispatus* human isolates are capable of growing on extracellular glycogen and we identified variation in a gene which correlated with this activity. The identified gene putatively encodes a pullulanase type I enzyme belonging to the glycoside hydrolase family 13 [38]. Its closest ortholog is an extracellular cell-attached pullulanase found in *L. acidophilus* [32]. The *L. crispatus* pullulanase gene described here carries three conserved domains, comprising an N-terminal carbohydrate-binding module family 41, a catalytic module belonging to the pullulanase super family and a C-terminal bacterial surface layer protein (SLAP) [39] (Figure 6). We observed that all except two of the strains in our study carry a copy of this gene. These two strains (RL31 and RL32), were no longer cultivable after their initial isolation. The six strains that showed less efficient or no growth on glycogen all showed variation in the N-terminal part of the pullulanase gene. All of these deletions are upstream of the carbohydrate-binding module in a sequence encoding a putative signal peptide. Furthermore, the presence of a SLAP-domain suggests that this enzyme is assigned to the outermost S-layer of the cell wall and is hence expected to be capable of degrading extracellular glycogen [32]. Further functional experiments are needed to fully characterize this pullulanase enzyme and to assess whether it degrades intra- or extracellular glycogen. Importantly, this pullulanase is likely part of a larger cluster of glycoproteins involved in glycogen metabolism in *L. crispatus*, which should be considered in future research.

**Figure 6.**
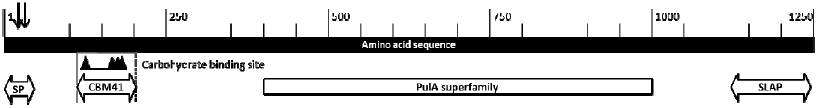
Schematic overview of the organization of the putative pullulanase type I encoding gene in *Lactobacillus crispatus*. The enzyme comprises three conserved domains including an N-terminal carbohydrate-binding module family 41 with specific carbohydrate binding sites, a catalytic module belonging to the pullulanase super family and a C-terminal bacterial surface layer protein (SLAP). The mutations (indicated by arrows) were located in an unconserved area that encodes a putative signal peptide (SP) that may be involved in subcellular localization. Abbreviations: SP: signal peptide; CBM41: carbohydrate-binding module family 41; PulA: pullulanase; SLAP: surface layer protein.

Of note, we analyzed just one *L. crispatus* strain per vaginal sample, while it is plausible that multiple strain types co-exist in the vagina. So strain variability in growth on glycogen (and other carbohydrates) might actually benefit the *L. crispatus* population as a whole and explain the variation in growth on glycogen that we observed, especially considering that glycogen availability may fluctuate along with oscillating estrogen levels during the menstrual cycle. When developing probiotics, it could thus be beneficial to select for *L. crispatus* strains that ferment different carbohydrates (in addition to glycogen) [8] and also to supplement the probiotic with a prebiotic [40, 41].

## Conclusion

Here we report whole-genome sequences of 28 *L. crispatus* human isolates. Our comparative study led to a total of three novel insights: 1) gene fragments encoding for a glycosyltransferase were disproportionally higher abundant among strains isolated from DVM, suggesting a role for cell surface glycoconjugates that shape vaginal microbiota-host interactions; 2) *L. crispatus* strains isolated from LVM do not differ from those isolated from DVM regarding the phenotypic traits studied here, including biofilm formation, pathogen inhibitory activity and carbohydrate utilization; and 3) *L. crispatus* is able to grow on glycogen and this correlates with the presence of a full-length pullulanase type I gene.

## METHODS

### L. crispatus strain selection

For this study, nurse-collected vaginal swabs were obtained from the Sexually Transmitted Infections clinic in Amsterdam, the Netherlands, from June to August 2012, as described previously by Dols *et al*. [4]. These vaginal samples came from women with LVM (Nugent score 0-3) and from women with DVM (Nugent score 7- 10). LVM and DVM vaginal swabs were plated on Trypton Soy Agar supplemented with 5% sheep serum, 0.25% lactic acid and pH set to 5.5 with acetic acid and incubated under microaerobic atmosphere (using an Anoxomat; Mart Microbiology B.V., the Netherlands) at 37°C for 48-72 hours. Candidate *Lactobacillus* spp. strains were selected based on colony morphology (white, small, smooth, circular, opaque colonies) and single colonies were subjected to 16S rRNA sequencing. One *L. crispatus* isolate per vaginal sample was taken forward for whole genome sequencing. A DNA library was prepared for these isolates using the Nextera XT DNA Library preparation kit and the genome was sequenced using the Illumina Miseq generate FASTQ workflow.

### Genome assembly and quality control

All analyses were run on a virtual machine running Ubuntu version 16.02. Contigs were assembled using the Spades assembly pipeline [42]. Contigs were discarded if they had less than 50% coverage with other assemblies or with the reference genome (N50 and NG50 values deviated more than 3 standard deviations from the mean as determined using QUAST [43]. The genomes were assembled with Spades 3.5.0 using default settings. The Spades pipeline integrates read-error correction, iterative k-mer (nucleotide sequences of length k) based short read assembling and mismatches correction. The quality of the assemblies was determined with Quast (History 2013) using default settings and the *Lactobacillus* crispatus ST1 strain as reference genome (Genbank FN692037).

### Genome annotation and comparative genome analysis

After assembly, the generated contigs were sorted with Mauve contig mover [44], using the *L. crispatus* ST1 strain as reference genome. Contaminating sequences of human origin and adaptor sequences were identified using BLAST and manually removed. The reordered genomes were annotated using the Prokka automated annotation pipeline [45] using default settings. Additionally, the genomes were uploaded to Genbank and annotated using the NCBI integrated Prokaryotic Genome Annotation Pipeline [46]. The annotated genomes were analyzed using the Sequence element enrichment analysis (SEER), which looks for an association between enriched k-mers and a certain phenotype [47]. Following the developer’s instructions, the genomes were split into k-mers using fsm-lite on standard settings and a minimum k-mer frequency of 2 and a maximum frequency of 28. The usage of k-mers enables the software to look for both SNPs as well as gene variation at the same time. After k-mer counting, the resulting file was split into 16 equal parts and g-zipped for parallelization purposes. In order to correct for the clonal population structure of bacteria, the population structure was estimated using Mash with default settings [48]. Using SEER, we looked for k-mers of various lengths that associated with whether the *L. crispatus* strains came from LVM or DVM. The results were filtered for k-mers with a chi-square test of association of <0.01 and a likelihood-ratio test p-value (a statistical test for the goodness of fit for two models) of <0.0001. The resulting list of k-mers was sorted by likelihood-ratio p and the top 50 hits were manually evaluated using BLASTx and BLASTn.

### Pan and accessory genome analysis

We used the bacterial pan genome analysis tool developed by Chaudhari *et al*. [49] using default settings. The circular image was created using CGview Comparison Tool [50] by running the build_blast_atlas_all_vs_all.sh script included in the package.

### Comparative phenotype experiments

Not all strains were (consistently) cultivable after their initial isolation, so experimental data was collected for a subset of the strains and could differ per experiment. The ratio of cultivable LVM and DVM strains was however similar for each experiment. For a full overview of experimental procedures, we refer to the Supplementary Information. In short, carbohydrate metabolism profiles were assessed using commercial API CH50 carbohydrate fermentation tests (bioMérieux, Inc., Marcy l’Etoile, France) according to the manufacturer’s protocol. To assess organic acid production, strains were grown on medium that mimicked vaginal secretions [26]. Total metabolite extracts from spent medium were assessed as previously described by Collins *et al*. [41]. Biofilm formation was assessed using the crystal violet assay as described by Santos *et al*. [51] and auto-aggregation as described by Younes et al. [52]. Antimicrobial activity against Neisseria gonorrhoeae was assessed by challenging *N. gonorrhoeae* (WHO-L strain) with varying (neutralized with NaOH to pH 7.0) dilutions of *L. crispatus* supernatants. Inhibitory effect was assessed as percentile difference in OD_600nm_ in a conditional stationary phase as compared to the control.

### Glycogen degradation assay

Starter cultures were grown in regular NYCIII glucose medium for 72 hours. For this assay, 1.1x carbohydrate deprived NYCIII medium was supplemented with water (negative control), 5% glucose (positive control) or 5% glycogen (Sigma-Aldrich, Saint Louis, US) and subsequently inoculated with 10% (v/v) bacterial culture (OD~0.5; 10^9^ CFU/ml). Growth on glycogen was compared to growth on NYCII without supplemented carbon source and to NYCIII with glucose. Growth curves were followed in a BioScreen (Labsystems, Helsinki, Finland). At least two independent experiments per strain were performed in triplicate.

## LIST OF ABBREVIATIONS

VM: vaginal microbiota
LVM: *Lactobacillus*-dominated vaginal microbiota
DVM: dysbiotic vaginal microbiota
COG: cluster ortholog genes
GT: glycosyltransferase
TSB: Trypton Soya Broth

## ETHICS APPROVAL AND CONSENT TO PARTICIPATE

The research proposed in this study was evaluated by the ethics review board of the Academic Medical Center (AMC), University of Amsterdam, The Netherlands. According to the review board no additional ethical approval was required for this study, as the vaginal samples used here were collected as part of routine procedure for cervical examinations at the STI clinic in Amsterdam (document reference number W12_086 # 12.17.0104). Clients of the STI clinic were notified that remainders of their samples could be used for scientific research, after anonymisation of client clinical data and samples. If the clients objected, their data and samples were discarded. This procedure has been approved by the AMC ethics review board (reference number W15_159 # 15.0193).

## CONSENT FOR PUBLICATION

Clients of the STI clinic were notified that remainders of their samples could be used for scientific research, after anonymisation of client clinical data and samples. If the clients objected, their data and samples were discarded. This procedure has been approved by the AMC ethics review board (reference number W15_159 # 15.0193).

## AVAILABILITY OF DATA AND MATERIAL

The 28 *Lactobacillus crispatus* sequenced genomes described in this paper have been deposited at DDBJ/ENA/GenBank under the accessions NKKQ00000000-NKLR00000000.

## AUTHORS’ CONTRIBUTIONS

RK, SB, HdV and FS conceptualized the study. CV and JS performed the experimental work, supervised by AdKA, SB and RK. JS performed the bio-informatic analyses, supervised by DW and RK. RH did the initial glycogen finding and provided further expertise. HT provided expertise for the glycosyltransferase finding and GR for the potential of probiotic applications. CV drafted the manuscript. All authors contributed to and approved the final manuscript.

## ACKNOWLEDGMENTS

We thank Dr. Titia Heijman of the Sexually Transmitted Infections clinic in Amsterdam, the Netherlands, for organizing the collection of the clinical vaginal samples. We thank Liesbeth Hoekman (TNO) for isolation and initial characterization of *Lactobacillus crispatus* strains. We thank Mark Sumarah and Justin Renaud for facilitating the metabolomics analysis. We also thank Dr. Zhi-Luo Deng for mining his transcriptomics data for the GT gene fragments and pullulanase gene.

